# Predictive Modeling of Acute Graft-versus-Host-Disease using Machine Learning on Immune Cell and Cytokine Profiles at Engraftment

**DOI:** 10.1101/2025.03.13.642957

**Authors:** Mohini Mendiratta, Praful Pandey, Shobhit Pandey, Sandeep Rai, Shuvadeep Ganguly, Archana Sasi, Ritu Gupta, Prabhat Singh Malik, Raja Pramanik, Sachin Kumar, Baibaswata Nayak, Riyaz Ahmed Mir, Sameer Bakhshi, Deepam Pushpam, Mukul Aggarwal, Aditya Kumar Gupta, Rishi Dhawan, Tulika Seth, Manoranjan Mahapatra, Ranjit Kumar Sahoo

## Abstract

**Background:** Acute Graft-versus-Host-Disease (aGvHD) is a major immune complication following allogenic hematopoietic stem cell transplantation (Allo-HSCT), initiated by conditioning regimen-associated tissue damage. It involves the complex interplay of immune cells and cytokines. Our study aims to leverage machine learning (ML) algorithms on the immune and cytokine profile of Allo-HSCT recipients to develop biomarker-based classification models to predict the onset of aGvHD at the time of engraftment.

**Materials and Methods:** Seventy patients diagnosed with hematological disorders who had undergone I^st^ Allo-HSCT were recruited from All India Institute of Medical Sciences, New Delhi, India. Peripheral blood (PB) was collected from the patients at the time of engraftment, and the immune cell subtypes and cytokine profiles were analyzed using flow cytometry and ELISA respectively. The individual cell counts were then processed using basic ML models, including support vector classifier with RBF kernel, Decision Tree, and Random Forest, chosen for their mathematical simplicity and feature importance advantage of Decision Trees and Random Forests. Various data settings were utilized in the study: combined immune and cytokine counts, immune cell counts only, cytokine counts only, T-cell counts only, NK cell counts only, dendritic cell counts only, and B-cell counts only. These configurations were selected to investigate how different data sets impact the prediction of aGvHD before its onset.

**Results:** At the engraftment flow cytometric analysis of reconstituted lymphocytes in patients who developed aGvHD revealed that there was a remarkable decrease in the ratio of CD4^+^/CD8^+^ T-cell and Tregs, with an increase in the cytotoxic regulatory NK-cell, dendritic cells, and B-cell. The levels of pro-inflammatory cytokines (IFN-γ, IL-1β, IP-10, TNF-α, IL-17α, IL-12p70, MIP-1α, MIP-1β, RANTES), and Th17-and Th1-cells were elevated with consequent decline of the levels of anti-inflammatory cytokine IL-10, IL-2, IL-4 and Th2-, Th9-cells. Machine learning based on 48 parameters [all immune cell subsets n=34 and all cytokines (n=14)]. The correlation heat map shows a higher correlation of aGvHD with the cytokine profile with or without immune cells (accuracy: 1), T-cell alone (accuracy: 0.96); NK-cell alone (accuracy: 0.93); dendritic cells alone (accuracy: 0.90), B-cell alone (accuracy: 0.86).

**Conclusion:** The current models classify perfectly, indicating the potential for a ML algorithm in predicting the onset of aGvHD. However, a study with a larger sample size is required to validate these classification models and mitigate the risk of overfitting observed due to the consistently high performance. The study also highlights the potential of cytokine profiles as a viable alternative to T-cell counts, as evidenced by the correlation heat map and classifier models. These findings provide valuable insights into dataset requirements and future directions for integrating ML models into aGvHD prediction.

## Introduction

The success of Allo-HSCT is significantly reliant on the intricate dynamics of immune cell reconstitution, particularly during the critical phase of neutrophil engraftment. Neutrophil engraftment is defined as the point after HSCT when the absolute neutrophil count (ANC) rises to ≥ 500 cells/μl and is maintained for three consecutive days without the need for growth factor support, indicating successful recovery of neutrophil production from the transplanted stem cells (1). It marks the initial phase of hematopoietic recovery, reduces the risk of infections, signifies successful donor stem cell engraftment (2), and represents a critical window for immune profiling, as immune reconstitution during this phase can influence the development of complications such as aGvHD. Furthermore, cytokines, as key mediators of immune regulation, play pivotal roles in modulating inflammatory responses (3,4), making them valuable biomarkers for immune dysregulation. A comprehensive understanding of the cellular composition including T-cell, NK cells, dendritic cells, and B-cell and their respective subtypes along with cytokine milieu during this phase is essential for comprehending the trajectory of immune recovery and the risk of aGvHD.

In recent years, the application of machine learning algorithms has emerged as a powerful approach for analyzing complex biological data, providing innovative means to develop predictive models for aGvHD based on the cellular and cytokine profiles observed during engraftment. This dual approach, combining immune and cytokine profiling with computational modeling, not only aims to improve early risk stratification for aGvHD but also provides novel insights into the immunopathogenesis of aGvHD. In addition to predictive modeling, we explored the kinetics of immune cell reconstitution in both aGvHD and non-aGvHD patients to elucidate distinct immune trajectories associated with aGvHD development.

To the best of our knowledge, this is the first study that comprehensively investigates the cellular (T-cell, NK cells, DCs, B-cell, and their subtypes) and cytokine profiles at the time of neutrophil engraftment in Allo-HSCT recipients and integration of high-dimensional immune profiling with advanced computational techniques, we applied ML algorithms to this dataset for the development of a predictive model for aGvHD.

## Materials and methods

### Study population

Patients diagnosed with hematological diseases who had undergone 1^st^ Allo-HSCT were recruited in the study. Patients (n=70) were recruited from the Department of Medical Oncology, Hematology, and Paediatrics Oncology, All India Institute of Medical Sciences, New Delhi between September 2020 and July 2023. Age and sex-matched healthy volunteers (n=20) were also included in the study.

Peripheral blood (PB) samples were collected from each patient at various time points-D+14, D+30, D+60, D+100, and D+180 following Allo-HSCT. In addition, if patients exhibited clinical signs and symptoms of aGVHD, blood samples were drawn at the onset of aGVHD.

### Enumeration of immune cellular subsets

White blood cells (WBCs) were isolated from PB using the bulk lysis method. Briefly, the PB sample was diluted with 1X RBC lysis buffer (Invitrogen, Thermo Fisher Scientific, USA) in a 1:3 ratio and incubated for 15 minutes at room temperature. Following incubation, the diluted blood was centrifuged at 2000 rpm for 5 minutes and the pellet was washed with 1X PBS (Thermo Fisher Scientific, USA) at 2000 rpm for 5 minutes. The resulting pellet was resuspended in 200μl 1X PBS (Thermo Fisher Scientific, USA) and the cells were stained with fluorochrome-conjugated anti-human monoclonal antibodies against surface markers including, CD27, CD20, CD19, CD141, KIR, CD16, CD1c, CD56, CD11c, CD123, CD7, HLA-DR, CD3, CD45RA, CD25, CD4, CD8 (Beckman Coulter, USA), IgD (Thermo Fisher Scientific, USA) for 40 minutes in dark at room temperature. Fluorochrome-conjugated anti-human monoclonal antibodies for chemokine receptors-CCR7, CXCR3, CCR10, CCR6, CCR4, and CXCR4 (Becton Dickinson, USA) were used to analyze subtypes of helper T-cell, cytotoxic T-cell, and effector memory helper T cell and their staining was performed at 37°C, allowing for the uptake of these markers from the cytoplasm to the cell surface. After surface staining, cells were fixed and permeabilized using BD Pharmingen™ Transcription Factor Buffer Set (Becton Dickinson, USA) followed by staining with anti-human monoclonal antibodies targeting cytoplasmic markers such as FOXP3 (Becton Dickinson, USA) for 30 minutes at room temperature in dark. A minimum of 5,000,000 events were acquired using the DxFlex flow cytometer (Beckman Coulter, USA) and the data was analyzed using the Kaluza software (Beckman Coulter, USA). A representation of immune cells and their subtypes, with profiling markers (mentioned in italics) used for immunophenotyping is depicted in Figure S1.

### Cytokines profiling of Allo-HSCT recipients

Cytokine levels, including IFN-γ, IL-10, MIP-1α, IL-1β, TNF-α, and IL-17α, were quantified using ELISA kits (Thermo Fisher Scientific, USA) according to the manufacturer’s instructions. Serum samples were obtained from three cohorts: healthy controls, non-aGvHD patients, and aGvHD patients, and the measurements in the patient cohorts were performed at the time of neutrophil engraftment, defined as an absolute neutrophil count (ANC) exceeding 500/μl.

### Machine learning algorithm for the development of prediction of aGvHD

The individual immune cell counts were analyzed using fundamental machine learning (ML) models, including a Support Vector Classifier (SVC) with a radial basis function (RBF) kernel, Decision Tree, and Random Forest algorithms. These models were specifically chosen for their unique strengths. The SVC with an RBF kernel excels at capturing non-linear relationships between features, making it a robust choice for complex biological datasets. Meanwhile, Decision Tree and Random Forest models offer interpretability through their ability to identify feature importance and mathematical simplicity and effectiveness in handling varied datasets. A range of data configurations was employed to investigate the impact of different immune parameters on the early prediction of aGvHD onset. These configurations were designed to assess the predictive power of distinct immune components, both individually and in combination. The first configuration combined immune cell counts and cytokine levels, offering a holistic perspective by integrating cellular and soluble immune mediators. The second configuration focused solely on immune cell counts, isolating the cellular component’s contribution to predictive accuracy. Similarly, the third configuration examined only cytokine levels to evaluate the role of soluble inflammatory mediators independently. Other configurations were tailored to investigate specific subsets of immune cells. For instance, T-cell counts were analyzed separately, given their critical role in aGVHD pathophysiology. Another configuration included both T-cell and NK-cell counts to explore the interplay between adaptive and innate lymphocyte populations. Dendritic cell counts were evaluated independently to understand the influence of antigen-presenting cells in predicting aGvHD. Lastly, B-cell counts were analyzed to assess the potential contribution of humoral immunity to disease onset. This systematic and comprehensive approach enabled a detailed understanding of how different immune components or their combinations contribute to the predictive modeling of aGVHD before its clinical manifestation.

### Statistical analysis

All statistical analyses were conducted using GraphPad Prism version 8.4.3. One-way and Tukey’s post hoc tests compared three or more groups. Data was shown as Mean±S.D. and a p-value of ≤ 0.05 was considered statistically significant.

## Results

### Patient characteristics

A total of 70 patients who underwent Allo-HSCT were included in this study. The most common underlying condition was AML, accounting for 35% (n = 25) of the cases, followed by aplastic anemia (22.8%, n = 16), thalassemia (12.8%, n = 9), ALL (11.4%, n = 8), MDS (4.28%, n = 3), CML-blast crisis (2.85%, n = 2), CLL (1.42%, n = 1), and other hematological disorders (8.5%, n = 6).

The majority of patients (70%, n = 49) underwent MSD transplantation, while 30% (n = 21) received haploidentical transplants. 47.14% (n = 33) of patients received ATG-based conditioning, 4.28% (n = 3) received PTCy alone, and 1.42% (n = 1) received a combination of ATG and PTCy. The remaining 47.14% (n = 33) underwent other chemotherapy-based conditioning regimens. TBI was administered to only one patient (1.42%), whereas the remaining 69 patients (98.58%) received only chemotherapy-based conditioning for immune ablation.

Grade II-IV aGvHD developed in 25 patients (35.71%) and 67.14% (n = 47) of recipients received CNI in combination with MTX, with or without corticosteroids, while 32.85% (n = 23) received CNI with MMF, with or without corticosteroids. The study design and patient recruitment flowchart are presented in Figure S2, and detailed clinical characteristics are summarized in Table S1.

### Aberrant T-cell reconstitution at the neutrophil engraftment contributes to the aGvHD progression in the later phase

T-cell are well-established mediators of aGvHD, playing a crucial role in APCs activation during the early phase of disease pathogenesis. In this study, we performed a comprehensive immunophenotypic analysis of T-cell, their subsets, including Th, Tc, and their respective naïve, effector, effector memory, and central memory populations, and Tregs in Allo-HSCT recipients at the time of neutrophil engraftment. The gating strategy used for their enumeration is depicted in Figure 1A-B.

**Figure 1:**
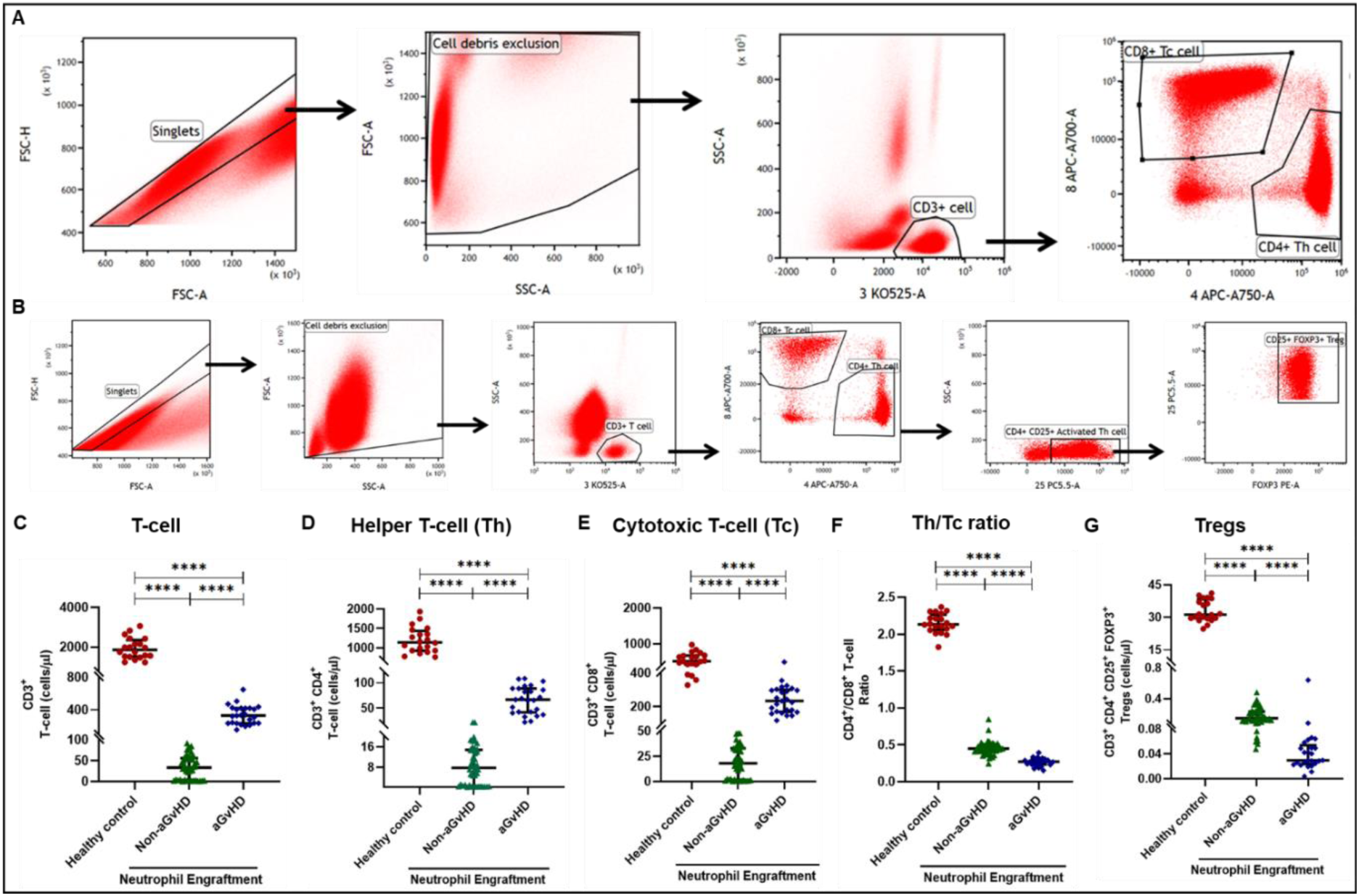
Profile of T-cell, helper T-cell, cytotoxic T-cell at the neutrophil engraftment of the patients who either did or did not develop aGvHD in the later phase and healthy control using flow cytometry. Dot plots illustrate the gating strategy for (A) T-cell, Th cell, and Tc cell. (B) Tregs. The scatter plot represents the absolute count (cells/μl) of (C) CD3^+^ T-cell. (D) CD3^+^ CD4^+^ T-cell. (E) CD3^+^ CD8^+^ T-cell. (F) CD4^+^/CD8^+^ T-cell ratio. (G) Tregs. Data are presented as the median with interquartile range for 20 healthy control and 70 patients. Statistical analysis: Mann-Whitney Test; ****≤0.0001. Abbreviations: Tregs: Regulatory Helper T-cell; aGvHD: Acute Graft-versus-Host-Disease

Our findings revealed that during the engraftment phase, all transplant recipients exhibit profound lymphopenia, characterized by significantly lower absolute cell counts compared to healthy controls, irrespective of whether they later developed aGvHD.

This state of lymphopenia results from the myeloablative conditioning regimen administered before transplantation. However, patients who subsequently developed aGvHD exhibited a markedly higher T-cell count at engraftment compared to those who did not (327.37 cells/µl vs 33.33 cells/µl; p ≤ 0.0001) (Figure 1C). This increase was evident in both Th (66.85 cells/µl vs 7.76 cells/µl; p ≤ 0.0001) (Figure 1D) and Tc populations (233.71 cells/µl vs 18.21 cells/µl; p ≤ 0.0001) (Figure 1E).

Interestingly, despite an overall increase in both Th and Tc subsets, aGvHD patients exhibited a significantly reduced Th/Tc ratio compared to non-aGvHD recipients (0.27 vs 0.45; p ≤ 0.0001) (Figure 1F), highlighting a predominant expansion of Tc cell over Th cell in this cohort. Additionally, aGvHD patients displayed a significantly lower Treg count compared to non-aGvHD recipients (0.030 vs 0.15; p ≤ 0.0001) (Figure 1G), indicative of a compromised immunoregulatory environment that may contribute to disease progression.

Furthermore, we analyzed all Th cell subsets (naïve, effector, effector memory, and central memory) in both patient cohorts at the time of neutrophil engraftment (Figure 2A). Patients who later developed aGvHD exhibited a significant increase in all Th subsets compared to those who did not (naïve: 6.58 cells/µl vs 0.06 cells/µl; p ≤ 0.0001, effector: 20.77 cells/µl vs 0.070 cells/µl; p ≤ 0.0001, effector memory: 26.30 cells/µl vs 0.88 cells/µl; p ≤ 0.0001, central memory: 1.34 cells/µl vs 0.07 cells/µl; p ≤ 0.0001) (Figure 2B-D). Interestingly, the effector Th cell population was significantly elevated in aGvHD patients compared to healthy controls (20.77 cells/µl vs 4.28 cells/µl; p ≤ 0.0001), whereas other Th subsets remained lower than in healthy controls, suggesting a selective expansion of effector Th cells in aGvHD.

**Figure 2:**
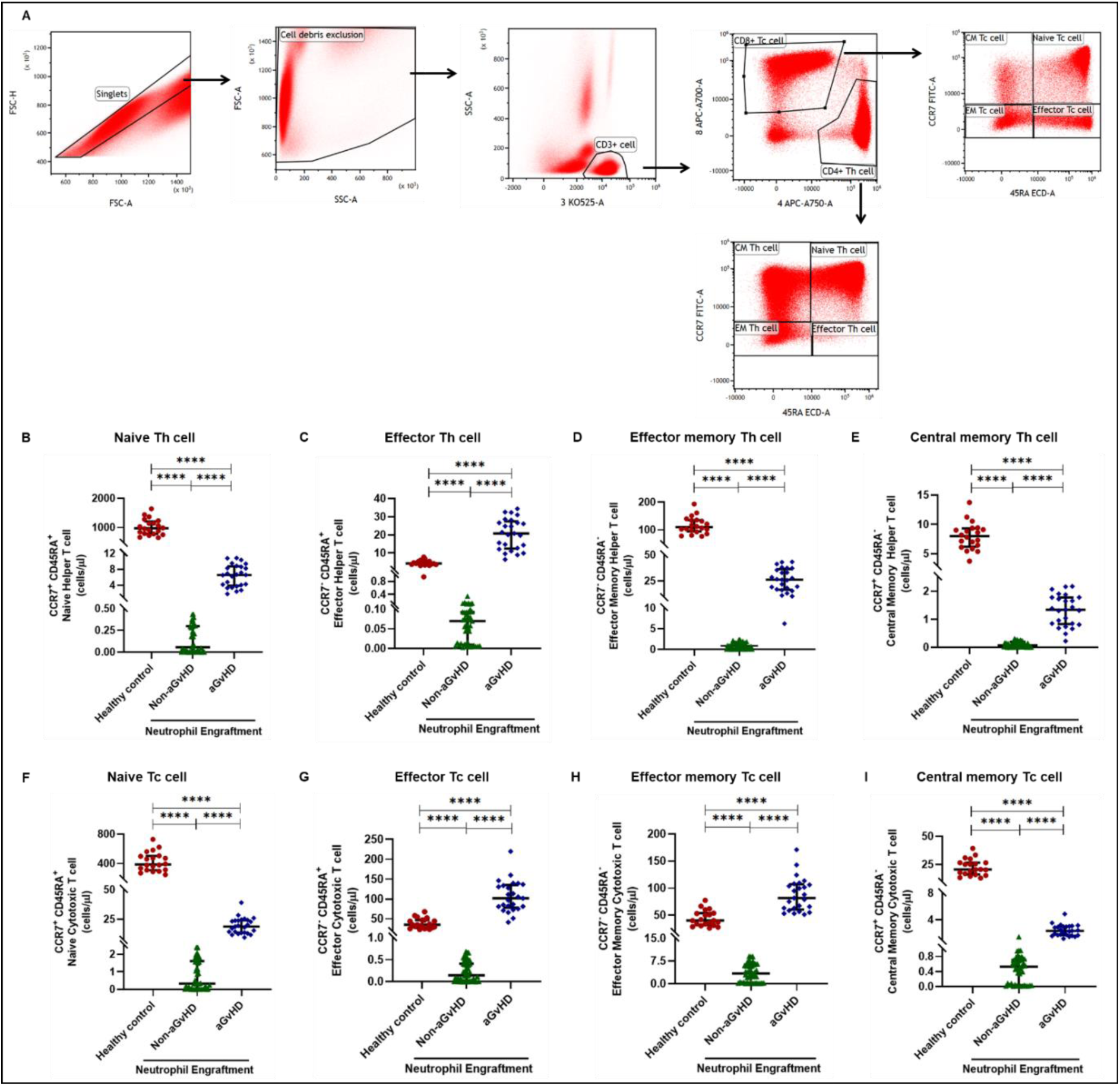
Profile of subtypes of Th and Tc at the neutrophil engraftment of the patients who either did or did not develop aGvHD in the later phase and healthy control using flow cytometry. Dot plots illustrate the gating strategy for (A) Subtypes of Th cell, and Tc cell. The scatter plot represents the absolute count (cells/μl) of (B) CCR7^+^ CD45RA^+^ Naïve Helper T-cell. (C) CCR7^-^ CD45RA^+^ Effector Helper T-cell. (D) CCR7^-^ CD45RA^-^ Effector Memory Helper T-cell. (E) CCR7^+^ CD45RA^-^ Central Memory Helper T-cell. (F) CCR7^+^ CD45RA^+^ Naïve Cytotoxic T-cell. (G) CCR7^-^ CD45RA^+^ Effector Cytotoxic T-cell. (H) CCR7^-^ CD45RA^-^ Effector Memory Cytotoxic T-cell. (I) CCR7^+^ CD45RA^-^ Central Memory Cytotoxic T-cell. Data are presented as the median with interquartile range for 20 healthy control and 70 patients (aGvHD=25; non-aGvHD=45). Statistical analysis: Mann-Whitney Test; ****≤0.0001. Abbreviations: Th: Helper T-cell; Tc: Cytotoxic T-cell; aGvHD: Acute Graft-versus-Host-Disease

Similarly, all Tc cell subsets (naïve, effector, effector memory, and central memory) were significantly increased in patients who developed aGvHD compared to those who did not (naïve: 18.70 cells/µl vs 0.33 cells/µl; p ≤ 0.0001, effector: 101.57 cells/µl vs 0.14 cells/µl; p ≤ 0.0001, effector memory: 81.80 cells/µl vs 3.38 cells/µl; p ≤ 0.0001, central memory: 2.34 cells/µl vs 0.53 cells/µl; p ≤ 0.0001) (Figure 2E-H). Notably, aGvHD patients exhibited a significantly higher proportion of both effector (101.57 cells/µl vs 35.11 cells/µl; p ≤ 0.0001) and effector memory Tc cell (81.80 cells/µl vs 40.16 cells/µl; p ≤ 0.0001) compared to healthy controls, highlighting a distinct expansion of these cytotoxic subsets in aGvHD pathogenesis.

Patients who later developed aGvHD exhibited a significantly higher proportion of Th1 (14.88 cells/μl vs 2.84 cells/μl; p ≤ 0.0001), Th17 (6.88 cells/μl vs 0.0003 cells/μl; p ≤ 0.0001), and Th22 (1.02 cells/μl vs 0.13 cells/μl; p ≤ 0.0001) cells, accompanied by a concurrent reduction in Th2 (0.14 cells/μl vs 1.16 cells/μl; p ≤ 0.05) and Th9 (0.09 cells/μl vs 0.15 cells/μl; p ≤ 0.05) subsets compared to those who did not develop aGvHD. Notably, Th17 cell were significantly elevated in the aGvHD patient cohort compared to healthy control (6.88 cells/μl vs 4.78 cells/μl; p ≤ 0.05), indicating a potential role for Th17 in aGvHD pathogenesis (Figure 3A-F).

**Figure 3:**
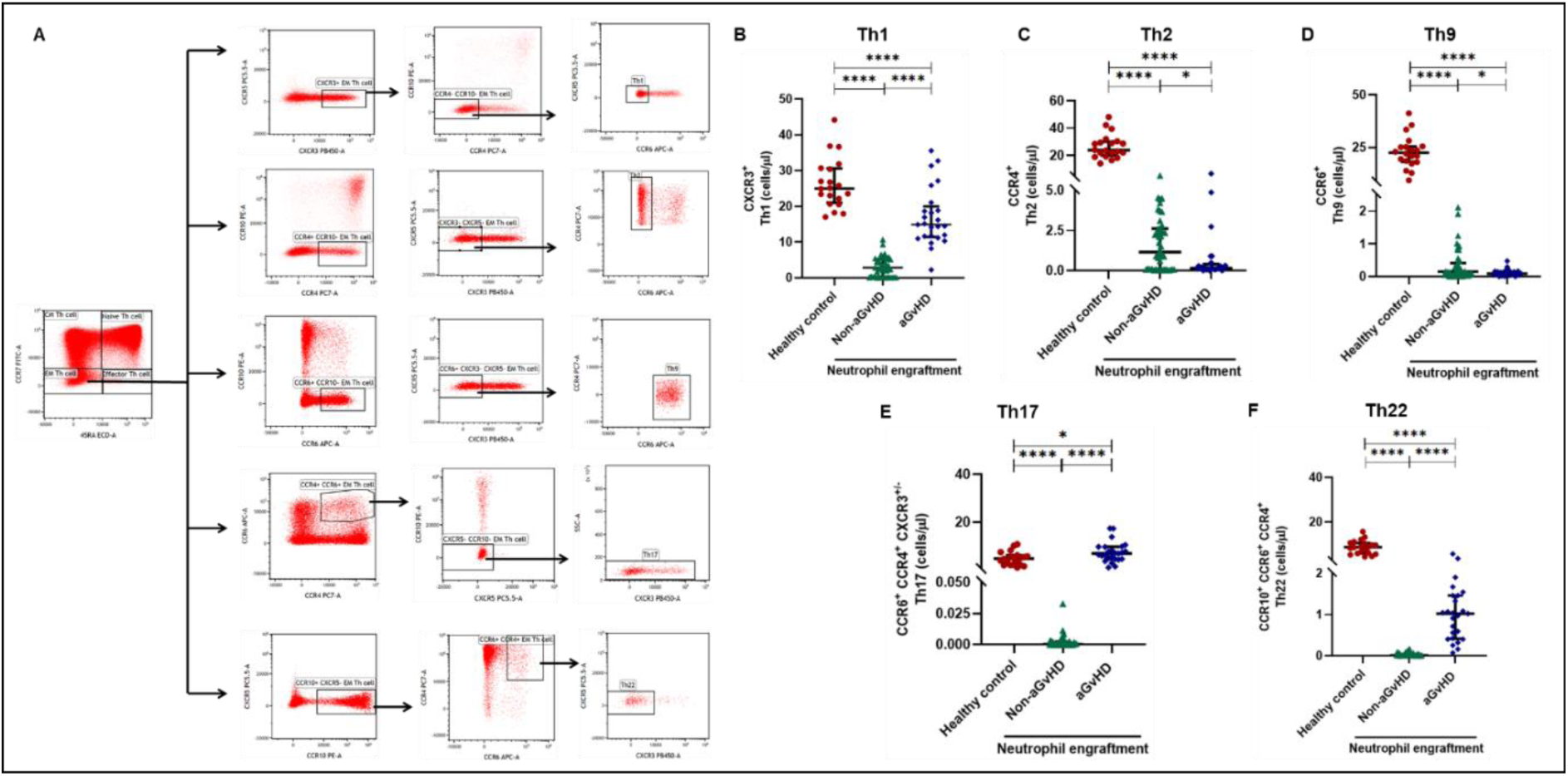
Profile of subtypes of effector memory Th at the neutrophil engraftment of the patients who either did or did not develop aGvHD in the later phase and healthy control using flow cytometry. Dot plots illustrate the gating strategy for (A) Subtypes of effector memory Th cell. The scatter plot represents the absolute count (cells/μl) of (B) Th1. (C) Th2. (D) Th9. (E) Th17. (F) Th22. Data are presented as the median with interquartile range for 20 healthy control and 70 patients (aGvHD=25; non-aGvHD=45). Statistical analysis: Mann-Whitney Test; *≤0.05; ****≤0.0001. Abbreviations: Th: Helper T-cell; aGvHD: Acute Graft-versus-Host-Disease

### Natural killer cell dynamics at the neutrophil engraftment associated with aGvHD onset

Natural killer cells play a critical role in aGvHD pathogenesis. In this study, we analyzed NK cells and their subsets in transplant recipients at the time of neutrophil engraftment to assess their potential in predicting aGvHD onset. The gating strategy used for NK cell enumeration and subtyping is depicted in Figure 4A.

**Figure 4:**
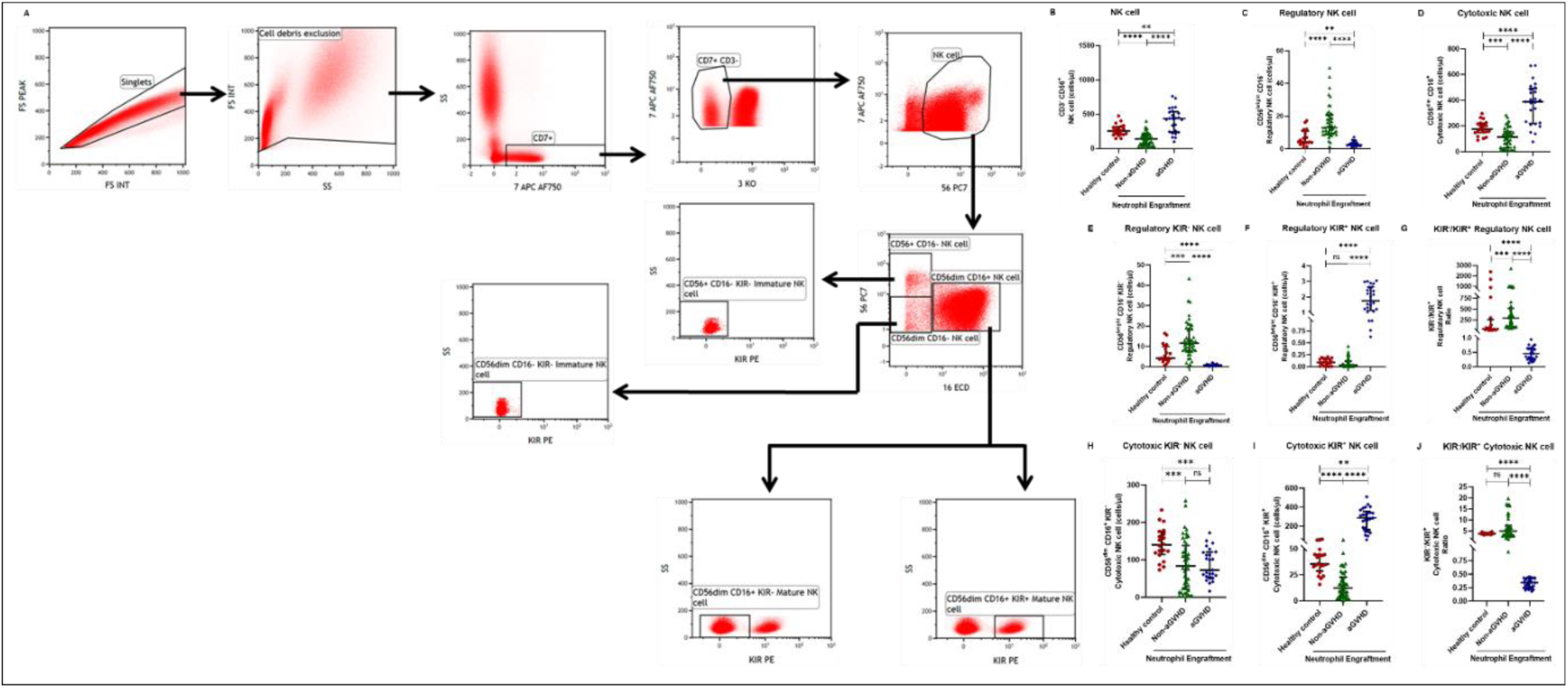
Natural killer cells and their subtypes at the neutrophil engraftment of the patients who either did or did not develop aGvHD in the later phase and healthy control using flow cytometry. Dot plots illustrate the gating strategy for (A) NK cell and their subtypes. The scatter plot represents the absolute count (cells/μl) of (B) CD3^-^ CD56^+^ NK cell. (C) CD56^bright^ CD16^-^ Regulatory NK cell. (D) CD56^dim^ CD16^+^ Cytotoxic NK cell (E) CD56^bright^ CD16^-^ KIR^-^ Regulatory NK cell. (F) CD56^dim^ CD16^+^ KIR^+^ Regulatory NK cell. (G) KIR^-^/KIR^+^ Regulatory NK cell. (H) CD56^dim^ CD16^+^ KIR^-^ Cytotoxic NK cell. (I) CD56^dim^ CD16^+^ KIR^+^ Cytotoxic NK cell. (J) KIR^-^/KIR^+^ Cytotoxic NK cell. Data are presented as the median with interquartile range for 20 healthy control and 70 patients (aGvHD=25; non-aGvHD=45). Statistical analysis: Mann-Whitney Test; *≤0.05; ****≤0.0001. Abbreviations: NK: Natural Killer Cell; KIR: Killer Immunoglobulin-like Receptors; aGvHD: Acute Graft-versus-Host-Disease

A comparative analysis of neutrophil engraftment profiles between patients who later developed aGvHD and those who did not revealed a significantly higher NK cell count in the aGvHD cohort (436.73 cells/μl vs 138.86 cells/μl; p ≤ 0.0001) (Figure 4B). This increase was primarily attributed to a significant expansion of cytotoxic NK cells (387.87 cells/μl vs 113.78 cells/μl; p ≤ 0.0001), accompanied by a concurrent reduction in regulatory NK cells (2.59 cells/μl vs 12.96 cells/μl; p ≤ 0.0001) in aGvHD patients compared to the non-aGvHD cohort (Figure 4C-D). Interestingly, NK cell counts were also significantly elevated in the aGvHD cohort compared to healthy controls (436.74 cells/μl vs 255.18 cells/μl; p ≤ 0.0001), with a similar trend observed for cytotoxic NK cells (387.87 cells/μl vs 174.73 cells/μl; p ≤ 0.0001).

Moreover, at the time of neutrophil engraftment, the aGvHD cohort exhibited a significantly higher fraction of regulatory NK cells expressing KIR (1.76 cells/μl vs. 0.03 cells/μl; p ≤ 0.0001), while the fraction of regulatory NK cells lacking KIR expression was markedly reduced (0.73 cells/μl vs 11.55 cells/μl; p ≤ 0.0001) compared to the non-aGvHD cohort (Figure 4E-F). This was reflected in a significant decrease in the KIR⁻/KIR⁺ regulatory NK cell ratio (0.46 vs 292.68; p ≤ 0.0001) in aGvHD patients (Figure 4G).

Notably, no significant difference was observed in the fraction of KIR⁻ cytotoxic NK cells between the aGvHD and non-aGvHD cohorts (73.11 cells/μl vs 84.10 cells/μl; p 0.6176). However, the fraction of KIR⁺ cytotoxic NK cells was significantly elevated in aGvHD patients (286.24 cells/μl vs 12.60 cells/μl; p ≤ 0.0001) (Figure 4H-I).

Consequently, the KIR⁻/KIR⁺ cytotoxic NK cell ratio was significantly reduced in the aGvHD cohort (0.34 vs 4.93; p ≤ 0.0001) (Figure 4J), suggesting a potential role of KIR⁺ cytotoxic NK cells in aGvHD pathogenesis.

### Elevated dendritic cells with the predominance of cDC2 drive aGvHD onset

Patients who developed aGvHD later exhibited significantly higher dendritic cell (DC) counts compared to both the non-aGvHD cohort (38.27 cells/µl vs 11.98 cells/µl; p ≤ 0.0001) and healthy controls (38.26 cells/µl vs 21.51 cells/µl; p ≤ 0.001) (Figure 5A-B). Notably, both pDC and cDC were significantly elevated in the aGvHD cohort relative to the non-aGvHD cohort (pDCs: 0.23 cells/µl vs 0.008 cells/µl; p ≤ 0.0001; cDCs: 31.25 cells/µl vs 5.05 cells/µl; p ≤ 0.0001) as well as healthy controls (pDCs: 0.23 cells/µl vs. 0.012 cells/µl; p ≤ 0.0001; cDCs: 31.25 cells/µl vs. 17.98 cells/µl; p ≤ 0.0001) (Figure 5C-D).

**Figure 5:**
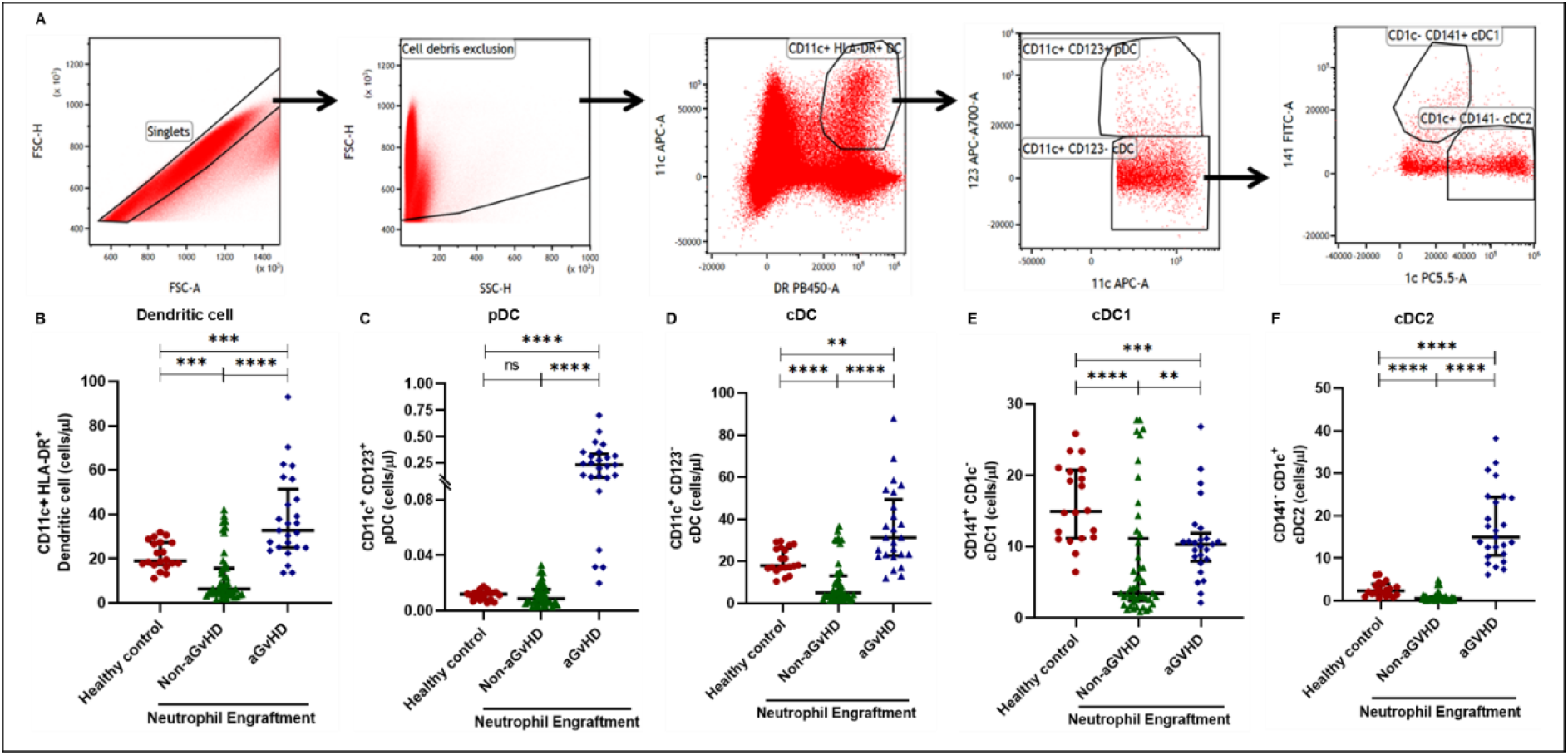
Dendritic cells and their subtypes at the neutrophil engraftment of the patients who either did or did not develop aGvHD in the later phase and healthy control using flow cytometry. Dot plots illustrate the gating strategy for (A) Dendritic cell and their subtypes. The scatter plot represents the absolute count (cells/μl) of (B) CD11c^+^ HLA-DR^+^ Dendritic cell. (C) CD11c^+^ CD123^+^ pDC. (D) CD11c^+^ CD123^-^ cDC (E) CD141^+^ CD1c^-^ cDC1. (F) CD141^-^ CD1c^+^ cDC2. Data are presented as the median with interquartile range for 20 healthy control and 70 patients (aGvHD=25; non-aGvHD=45). Statistical analysis: Mann-Whitney Test; **≤0.01; ***≤0.001; ****≤0.0001. Abbreviations: pDC: Plasmacytoid Dendritic Cell; cDC: Conventional Dendritic Cell; aGvHD: Acute Graft-versus-Host-Disease

Within the cDC subset, both cDC1 and cDC2 fractions were elevated in aGvHD compared to the non-aGvHD cohort (cDC1: 10.31 cells/µl vs 3.46 cells/µl; p ≤ 0.01; cDC2: 14.98 cells/µl vs. 0.55 cells/µl; p ≤ 0.0001). Furthermore, cDC2 was significantly more abundant in the aGvHD cohort than in healthy controls (14.98 cells/µl vs. 2.35 cells/µl; p ≤ 0.0001) (Figure 5E-F).

### Elevated B-cell count at the neutrophil engraftment predicted aGVHD onset in the transplant recipients

In addition to T-cell, NK cells, and dendritic cells, B-cell and their subtypes were assessed in Allo-HSCT recipients at the time of neutrophil engraftment. The gating strategy for B-cell enumeration and subtyping is depicted in Figure 6A. Our analysis revealed that transplant recipients were lymphopenic at neutrophil engraftment. However, the absolute B-cell count was significantly higher in patients who later developed aGvHD compared to those in the non-aGvHD cohort (18.68 cells/µl vs 0.52 cells/µl; p ≤ 0.0001) (Figure 6B). Similarly, all B-cell subtypes, including naïve B-cell (9.39 cells/µl vs 0.31 cells/µl; p ≤ 0.0001), USM B-cell (0.25 cells/µl vs. 0.020 cells/µl; p ≤ 0.0001), SM B-cell (0.43 cells/µl vs. 0.037 cells/µl; p ≤ 0.0001), and DNSM B-cell (2.51 cells/µl vs. 0.069 cells/µl; p ≤ 0.0001), were significantly elevated in the aGvHD cohort compared to non-aGvHD recipients (Figure 6C–F).

**Figure 6:**
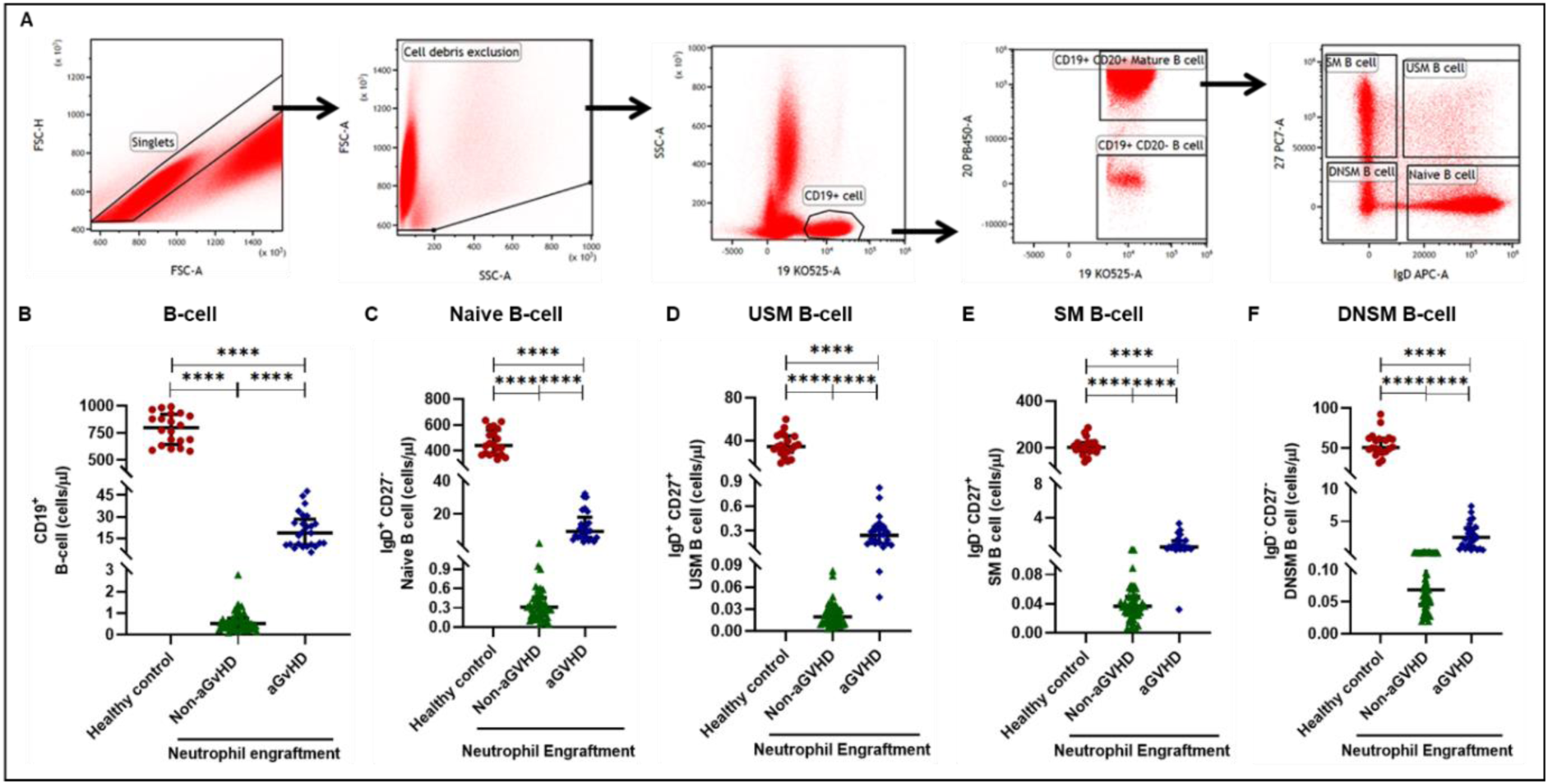
B-cell and their subtypes at the neutrophil engraftment of the patients who either did or did not develop aGvHD in the later phase and healthy control using flow cytometry. Dot plots illustrate the gating strategy for (A) B-cell and their subtypes. The scatter plot represents the absolute count (cells/μl) of (B) CD19^+^ B-cell. (C) IgD^+^ CD27^-^ Naïve B-cell. (D) IgD^+^ CD27^+^ USM B-cell. (E) IgD^-^ CD27^+^ SM B-cell. (F) IgD^-^ CD27^-^ DNSM B-cell. Data are presented as the median with interquartile range for 20 healthy control and 70 patients (aGvHD=25; non-aGvHD=45). Statistical analysis: Mann-Whitney Test; ****≤0.0001. Abbreviations: USM: Unswitched Memory; SM: Switched Memory; DNSM: Double Negative Switched Memory; aGvHD: Acute Graft-versus-Host-Disease

### Imbalance in cytokines at the neutrophil engraftment predicted aGvHD onset in the transplant recipients

In addition to cellular profiling, cytokines and chemokines were quantified in transplant recipients at the time of neutrophil engraftment to assess their correlation with aGVHD occurrence. Our analysis revealed that patients who later developed aGVHD exhibited significantly higher levels of pro-inflammatory cytokines, including IFN-γ, IL-1β, IP-10, TNF-α, IL-17A, IL-12 (p70), IL-6, IL-5, RANTES, MIP-1α, and MIP-1β, along with lower levels of anti-inflammatory cytokines such as IL-2, IL-4, and IL-10 (Figure 7A–N). This cytokine profile, characterized by an elevation in pro-inflammatory mediators and a concurrent reduction in anti-inflammatory cytokines, was observed in aGVHD patients compared to healthy controls and transplant recipients who did not develop aGVHD in the later post-transplant phase.

**Figure 7:**
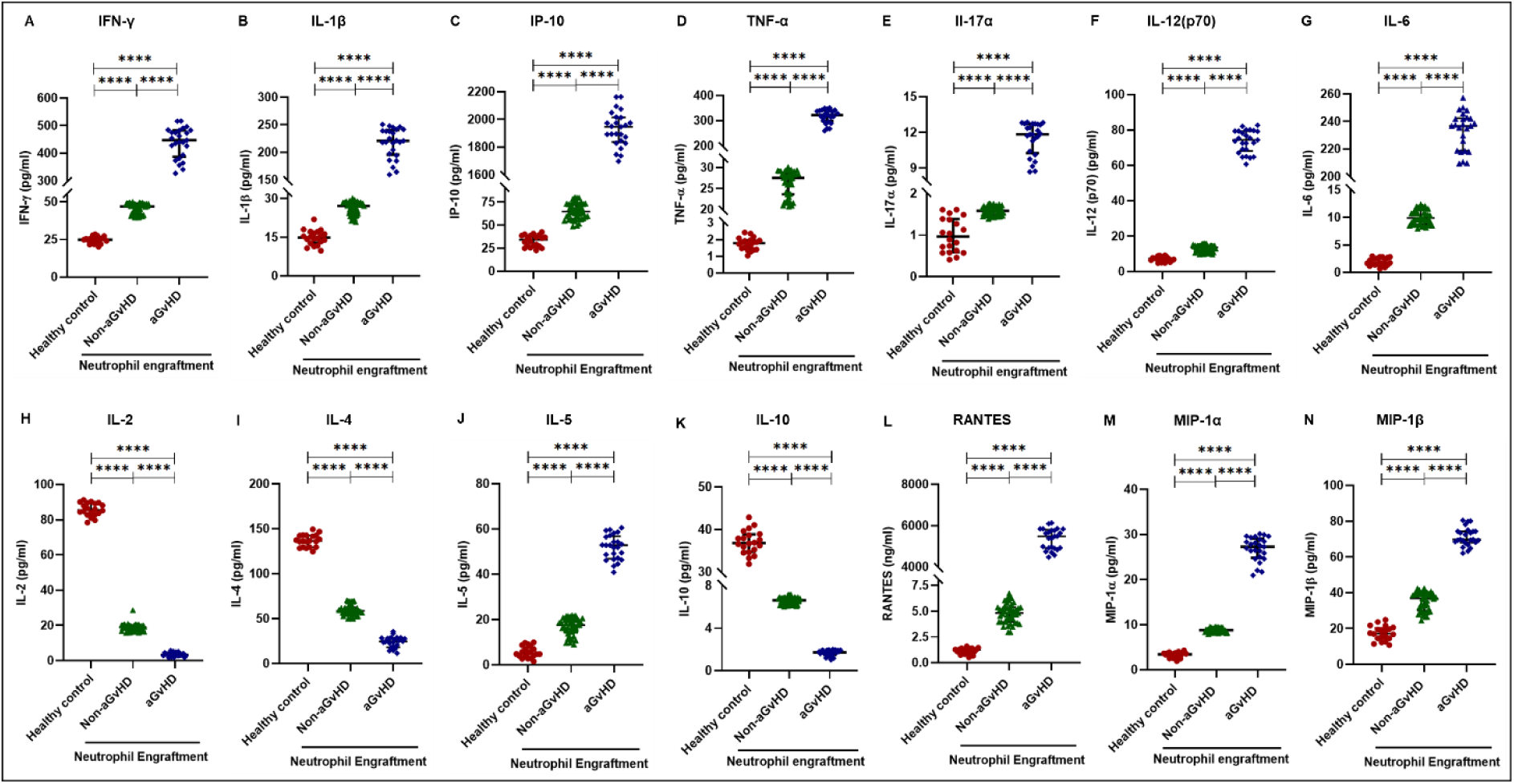
Cytokines and chemokines profile at the neutrophil engraftment of the patients who either did or did not develop aGvHD in the later phase and healthy control. The scatter plot represents the concentration (pg/ml) of (A) IFN-γ. (B) IL-1β. (C) IP-10 (ng/ml). (D) TNF-α (E) IL-17α. (F) IL-12 (p70). (G) IL-6. (H) IL-2. (I) IL-4. (J) IL-5. (K) IL-10. (L) RANTES. (M) MIP-1α. (N) MIP-1β. Data are presented as the median with interquartile range for 20 healthy control and 70 patients (aGvHD=25; non-aGvHD=45). Statistical analysis: Mann-Whitney Test; ****≤0.0001. Abbreviations: IFN-γ: Interferon-γ; IL-: Interleukin; MIP: Macrophage Inflammatory Protein; RANTES: Regulated on Activation, Normal T-cell Expressed and Secreted; aGvHD: Acute Graft-versus-Host-Disease

### Machine learning (ML) algorithm identified cytokine levels as better predictive markers than cellular parameters at the time of neutrophil engraftment

Our ML analysis of 48 immune parameters (34 immune cell subsets and 14 cytokines) revealed distinct immunological dynamics at the time of neutrophil engraftment, which may serve as potential biomarkers for aGvHD prediction. Due to the complexity and high-dimensional nature of these immune variables, we employed a machine learning approach to integrate and analyze the dataset, enabling the development of predictive models for early risk stratification and personalized therapeutic interventions (Figure 8A).

**Figure 8:**
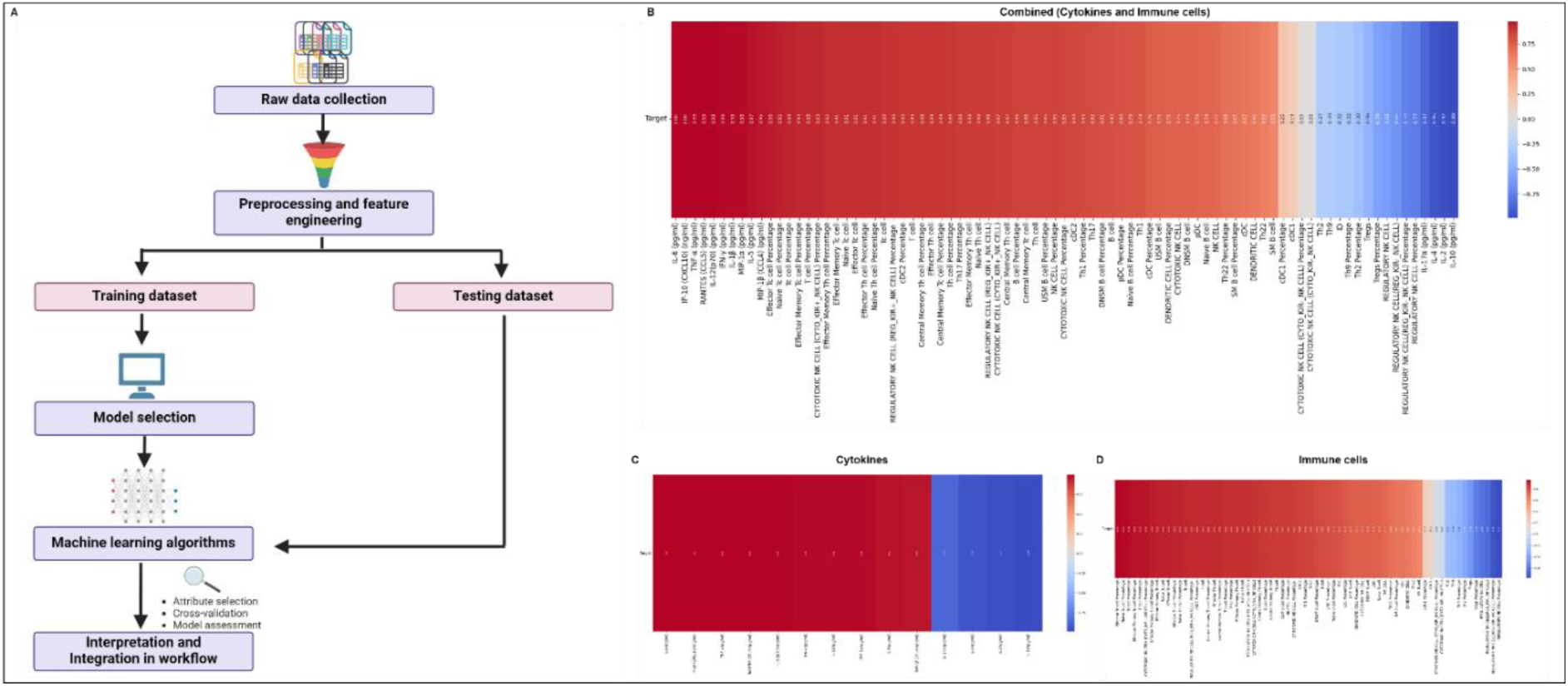
Integration of the ML algorithm in developing a predictive model for aGvHD using the cellular and cytokine profiles of transplant recipients at neutrophil engraftment. (A) Schematic representation of the methodology adopted for the ML approach. The correlation map shows (A) A combination of cytokines and cellular profiles. (B) Cytokine profile only. (C) Cellular profile only of transplant recipients at neutrophil engraftment for the identification of predictive marker for aGvHD.

The initial application of the machine learning model to combined immune cell and cytokine data demonstrated that cytokines at the neutrophil engraftment exhibited a stronger correlation with aGvHD onset in the later phase post-transplant compared to immune cell subsets, as depicted in the heat map (Figure 8B). Further analysis of cytokine correlations (Figure 8C) identified a highly positive correlation (1.00) with IL-6 and IP-10, followed by strong positive correlations with TNF-α (0.99), RANTES (0.99), IL-12 (p70) (0.99), and IFN-γ (0.99), IL-1β and MIP-1α (0.98), IL-5 (0.97), and MIP-1β (0.96). Conversely, a negative correlation was observed with IL-17A (-0.87), IL-4 (-0.95), IL-2 (-0.97), and IL-10 (-0.99).

The immune cell correlation heat map (Figure 8D, 9A-D) revealed a strong positive correlation (ranging from 0.96 to 0.59) between aGvHD and multiple immune subsets, including total T-cell, helper T-cell, cytotoxic T-cell, NK cells, dendritic cells, B-cell, and some of their subpopulations. A weaker correlation was observed with cDC1 (0.25), while no correlation was detected with KIR⁻ cytotoxic NK cells (0.00). In contrast, a negative correlation was identified with Th9 (-0.32), Th2 (-0.38), Tregs (-0.58), regulatory NK cells (-0.77), and KIR⁻ regulatory NK cells (-0.74).

The cellular and cytokine profiling of patients at the neutrophil engraftment who did develop or did not develop aGvHD in the later phase post-transplant was visualized using Uniform Manifold Approximation and Projection (UMAP), as shown in Figure 9E-I. UMAP analysis revealed distinct clustering patterns between the two groups, highlighting immunological differences at the time of neutrophil engraftment. Notably, cytokine profiles (Figure 9E) exhibited a more pronounced separation between aGvHD and non-aGvHD groups, suggesting their potential as robust discriminative markers. Similarly, immune cell distribution (Figure 9F-I) demonstrated a degree of separation, with certain cell subsets displaying differential abundance across groups.

**Figure 9:**
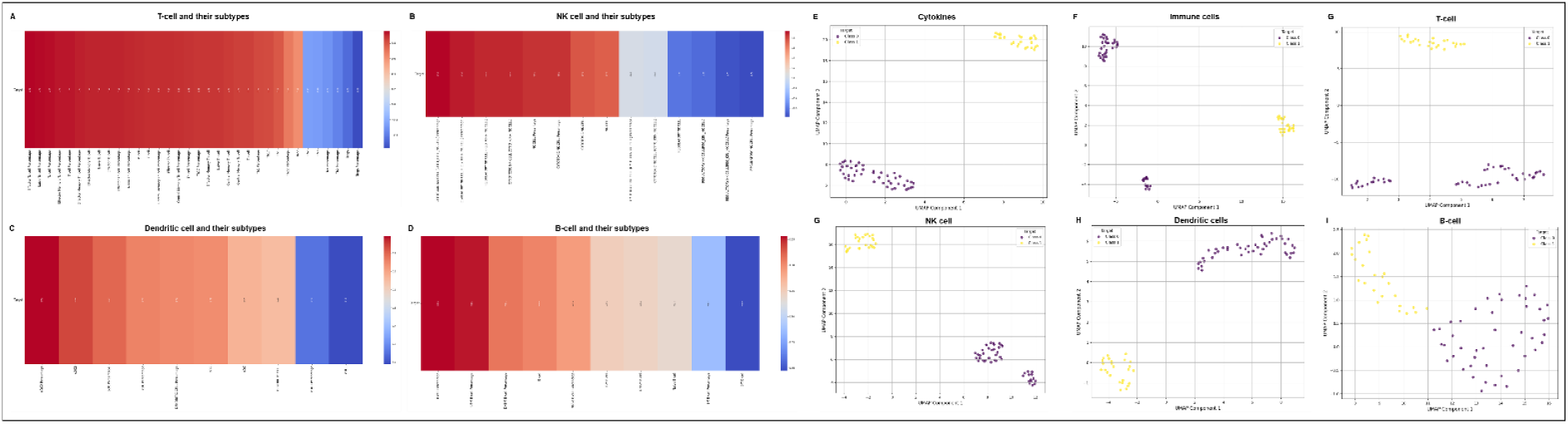
The correlation map shows (A) T-cell and their subtypes. (B) NK cells and their subtypes. (C) Dendritic cells and their subtypes (D) B-cell and their subtypes. The UMAP shows (E) Cytokines. (F) Immune Cells. (G)T-cell. (H) NK cells. (I) Dendritic cells. (J) B-cell of transplant recipients at neutrophil engraftment for the identification of predictive marker for aGvHD. Abbreviations: NK: Natural Killer

### Distinct immune reconstitution associated with aGvHD and non-aGvHD patients

In our study, we evaluated the kinetics of immune cell reconstitution in aGvHD and non-aGvHD patients from the point of neutrophil engraftment D+14 up to D+180. Longitudinal assessments were performed at multiple time points (D+30, +60, +100, and +180) to characterize differences in cellular reconstitution dynamics between the two groups.

Our findings revealed a significant difference in the pattern of T-cell reconstitution between aGvHD and non-aGvHD patients from D+14 to D+100. aGvHD patients exhibited a higher absolute T-cell count throughout this period compared to the non-aGvHD cohort, suggesting a distinct immune reconstitution trajectory associated with GvHD development. However, by D+180, the T-cell counts between the two groups converged, with no statistically significant difference observed (Figure 10A).

**Figure 10:**
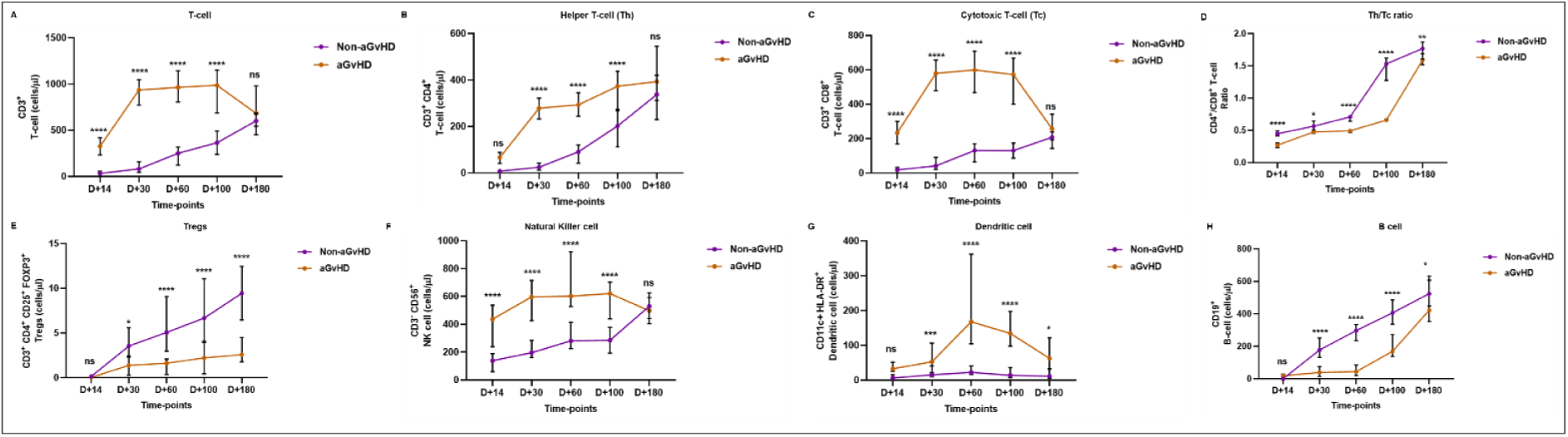
Kinetics of immune reconstitution of aGvHD and non-aGvHD patients. The line graphs represent the absolute count (cells/μl) from D+14 to D+180 of (A) CD3^+^ T-cell. (B) CD3^+^ CD4^+^ T-cell. (C) CD3^+^ CD8^+^ T-cell. (D). CD4^+^/CD8^+^T-cell ratio. (E) CD3^+^ CD4^+^ CD25^+^ FOXP3^+^ Tregs. (F) CD3^-^ CD56^+^ NK cell. (G) CD11c^+^ HLA-DR^+^ Dendritic cells. (H) CD19^+^ B-cell. Data are presented as the median with interquartile range for aGVHD patients (n=25) and non-aGvHD patients (n=45). Statistical analysis: Mann-Whitney Test; *≤0.s05; **≤0.01; ***≤0.001; ****≤0.0001. Abbreviations: NK: Natural Killer; Tregs: Regulatory Helper T-cell; aGvHD: Acute Graft-versus-Host-Disease

Similarly, aGvHD patients exhibited significantly higher counts of both CD4⁺ and CD8⁺ T-cell from D+14 to D+100 compared to the non-aGvHD cohort, indicating a differential T-cell reconstitution pattern associated with aGvHD development. Notably, by D+180, these differences were no longer observed. The convergence in CD4⁺ T-cell counts was driven by an increase in CD4⁺ T-cell in non-aGvHD patients, reaching levels comparable to those observed in the aGvHD cohort. In contrast, the CD8⁺ T-cell count in aGvHD patients declined over time, eventually aligning with the levels seen in non-aGvHD patients by D+180 (Figure 10B, C).

A similar trend was observed in the CD4⁺/CD8⁺ T-cell ratio, which exhibited a significant difference between the two groups. In the non-aGvHD cohort, the CD4⁺/CD8⁺ T-cell ratio remained consistently higher than that of the aGvHD cohort until D+100, after which it reached a plateau. In contrast, the aGVHD cohort demonstrated a substantial increase in the CD4⁺/CD8⁺ ratio from D+100 to D+180, eventually approximating the levels observed in the non-aGVHD cohort. This progressive rise in the CD4⁺/CD8⁺ ratio suggests a dynamic shift in T-cell composition, likely indicative of immune reconstitution or regulatory mechanisms influencing T-cell homeostasis in aGvHD patients (Figure 10D).

At D+14, no significant difference was observed in the absolute Tregs count between the aGvHD and non-aGvHD cohorts. However, from D+30 to D+180, both groups exhibited a significant increase in Tregs count. Notably, throughout the follow-up period, the Tregs count remained lower in the aGvHD cohort compared to the non-aGvHD cohort. By D+180, a marked increase in Tregs count was observed in the non-aGvHD cohort between D+100 and D+180, whereas in the aGvHD cohort, Treg levels reached a plateau between D+100 and D+180 (Figure 10E).

In addition to the T-cell compartment, the kinetics of NK cells, dendritic cells, and B-cell were assessed in both cohorts. A significant difference in NK cell count was observed between the groups, with the aGvHD cohort exhibiting a significantly higher NK cell count from D+14 to D+100 compared to the non-aGvHD cohort. In the aGvHD group, NK cell levels increased from D+14 to D+30, followed by a plateau from D+30 to D+100, and a subsequent decline from D+100 to D+180. In contrast, the non-aGvHD cohort showed a steady increase in NK cell count from D+14 to D+60, followed by a plateau until D+100, and a marked increase from D+100 to D+180. By D+180, no significant difference in NK cell count was observed between the cohorts (Figure 10F). The reconstitution dynamics of dendritic cells in the aGvHD cohort were highly variable. While there was no significant difference in their absolute count at D+14, a subsequent increase was observed at D+30 in both cohorts. In the aGvHD group, a pronounced expansion occurred from D+30 to D+60, followed by a decline from D+60 to D+180. In contrast, the non-aGvHD cohort exhibited a modest increase from D+30 to D+60, followed by a slight decline from D+60 to D+100, and a stabilization from D+100 to D+180. Notably, the absolute dendritic cell count remained higher in the aGvHD cohort throughout the follow-up period (Figure 10G).

For B-cell, no significant difference was observed between the cohorts at D+14. In the aGvHD group, B-cell counts remained similar from D+14 to D+60, followed by a substantial increase from D+60 to D+180. In the non-aGvHD cohort, a continuous and pronounced increase in B-cell count was observed from D+14 to D+180. Despite this expansion, B-cell counts remained lower in the aGvHD cohort compared to the non-aGvHD cohort throughout the follow-up period (Figure 10H).

## Discussion

We investigated the immunological dynamics at the time of neutrophil engraftment in patients who underwent Allo-HSCT. Our findings provide significant insights into the cellular and cytokine profiles associated with aGvHD, highlighting the critical role of T-cell, NK cells, dendritic cells, and B-cell in modulating post-transplant outcomes.

The majority of our cohort consisted of patients with AML and aplastic anemia, reflecting the predominance of these disorders in transplant populations. Notably, aGvHD developed in a substantial percentage of recipients, underscoring its importance as a major complication in Allo-HSCT, with previous studies indicating that nearly 30%-50% of patients may experience some degree of aGvHD after transplantation (5,6). Our research elucidated those patients who progressed to aGvHD exhibited significantly elevated T-cell counts during the engraftment phase, particularly in both Th and Tc subsets, corroborating earlier findings that highlighted the role of T-cell expansion in the pathophysiology of aGvHD (7).

The results demonstrated a marked increase in the Th1 and Th17 populations among aGvHD patients, consistent with the recognized pro-inflammatory roles of these subsets in promoting alloreactivity post-transplantation (8–11). In contrast, the reduction in the Th2 and Treg populations observed in aGvHD patients reflects a compromised immunoregulatory environment, as Tregs are crucial for maintaining tolerance and mitigating hyper-reactive immune responses (12,13). This dynamic imbalance may facilitate the onset of aGvHD, as Tregs provide essential control over the activity of potentially harmful effector T-cell (14,15).

Interestingly, the higher NK cell counts and the distinct increase in cytotoxic NK cells in aGvHD patients reinforce the notion that NK cells are significant contributors to graft-versus-host reactions. The prominent elevation of KIR^+^ cytotoxic NK cells observed in our findings mirrors previous reports that have suggested that these cells participate in sustaining inflammatory responses and enhancing tissue damage in aGvHD through direct cytotoxic effects on host tissues (Fei Gao et al, 2020).

Additionally, our analysis of dendritic cells revealed significantly increased counts in aGvHD patients, which is pivotal given that dendritic cells serve as powerful antigen-presenting cells that can activate T-cell responses. The deleterious role of dendritic cells in aGvHD development has been previously documented, indicating that their activation can lead to the amplification of inflammatory responses vital to the pathogenesis of GvHD (16,17).

Our study revealed that elevated levels of pro-inflammatory cytokines alongside reduction in anti-inflammatory cytokines further emphasizes the skewed balance of the immune milieu towards inflammatory pathways in aGvHD patients. This finding follows studies illustrating how alterations in the cytokine landscape facilitate allogeneic immune responses (3,18). Our machine learning model reinforced the association between specific cytokine profiles and aGvHD onset, demonstrating the promising potential for utilizing these immune parameters in predictive models that could guide personalized therapeutic vigilance post-transplant.

The delayed reconstitution of Tregs in the aGvHD cohort emphasizes the significance of an intact regulatory environment for preventing alloreactive responses. Studies have shown that restoring levels of Tregs can alleviate the severity of aGvHD, underscoring the need for therapeutic strategies that focus on enhancing Tregs function or expansion in the post-transplant period (19,20). This is crucial when considering the risk of infection due to immune suppression, as seen with aGvHD management regimens that utilize immunosuppressive therapies that may hinder Treg recovery.

## Conclusion

Our comprehensive analysis underscores the critical immunological profiles that emerge at the time of neutrophil engraftment and their implications for the prediction of aGVHD development. The correlation of cellular and cytokine responses with existing literature, we emphasize the complex interplay between different immune subsets and the necessity for refined monitoring and potential therapeutic interventions aimed at optimizing graft acceptance and reducing aGvHD incidents.

## Author’s Contributions

MM performed the experiments, acquired and analyzed data, interpreted the results, and wrote the manuscript. PP, SP, SR, SG, AS, RG, PSM, RP, SK, BN, and RAM contributed to data interpretation and analysis. SM provided resources, conceptualized the study, and designed and supervised the experiments. SB, DP, MA, AKG, RD, TS, MM provided patient samples and their clinical details. RKS provided funding and resources, conceptualized the study, designed and supervised the experiments, analyzed data, and edited the manuscript. All authors critically reviewed and approved the final version of the manuscript.

## Data availability

The data that support the findings of this study will be made available from the corresponding author upon reasonable request.

## Acknowledgment

The authors express their gratitude to the All India Institute of Medical Sciences (AIIMS), New Delhi, India for facilitating the execution of the study.

## Abbreviations

Allo-HSCT: Allogeneic Hematopoietic Stem Cell Transplantation
aGvHD: Acute Graft-versus-Host-Disease
ANC: Absolute Neutrophil Count
ML: Machine Learning
ELISA: Enzyme Linked Immunosorbent Assay
PB: Peripheral Blood
Tregs: Regulatory Helper T-cell
NK: Natural Killer
IFN-γ: Interferon-gamma
IL-1β: Interleukin-1beta
IP-10: Interferon Gamma-Induced Protein 10
TNF-α: Tumor Necrosis Factor-alpha
IL-17α: Interleukin-17 alpha
IL-12p70: Interleukin-12p70
MIP-1α: Macrophage Inflammatory Protein 1-alpha
MIP-1β: Macrophage Inflammatory Protein 1-beta
RANTES: Regulated Upon Activation, Normal T-cell Expressed and Secreted
SVC: Support Vector Classifier
RBF: Radial Basis Function
AML: Acute Myeloid Leukemia
ALL: Acute Lymphoblastic Leukemia
CML: Chronic Myeloid Leukemia
MDS: Myelodysplastic Syndrome
CLL: Chronic Lymphocytic Leukemia
MSD: Myelodysplastic Syndromes
PTCy: Post Transplant Cyclophosphamide
ATG: Antithymocyte globulin
TBI: Total Body Irradiation
MTX: Methotrexate
CNI: Calcineurin Inhibitors
MMF: Mycophenolate mofetil

---

**Figure S1:**
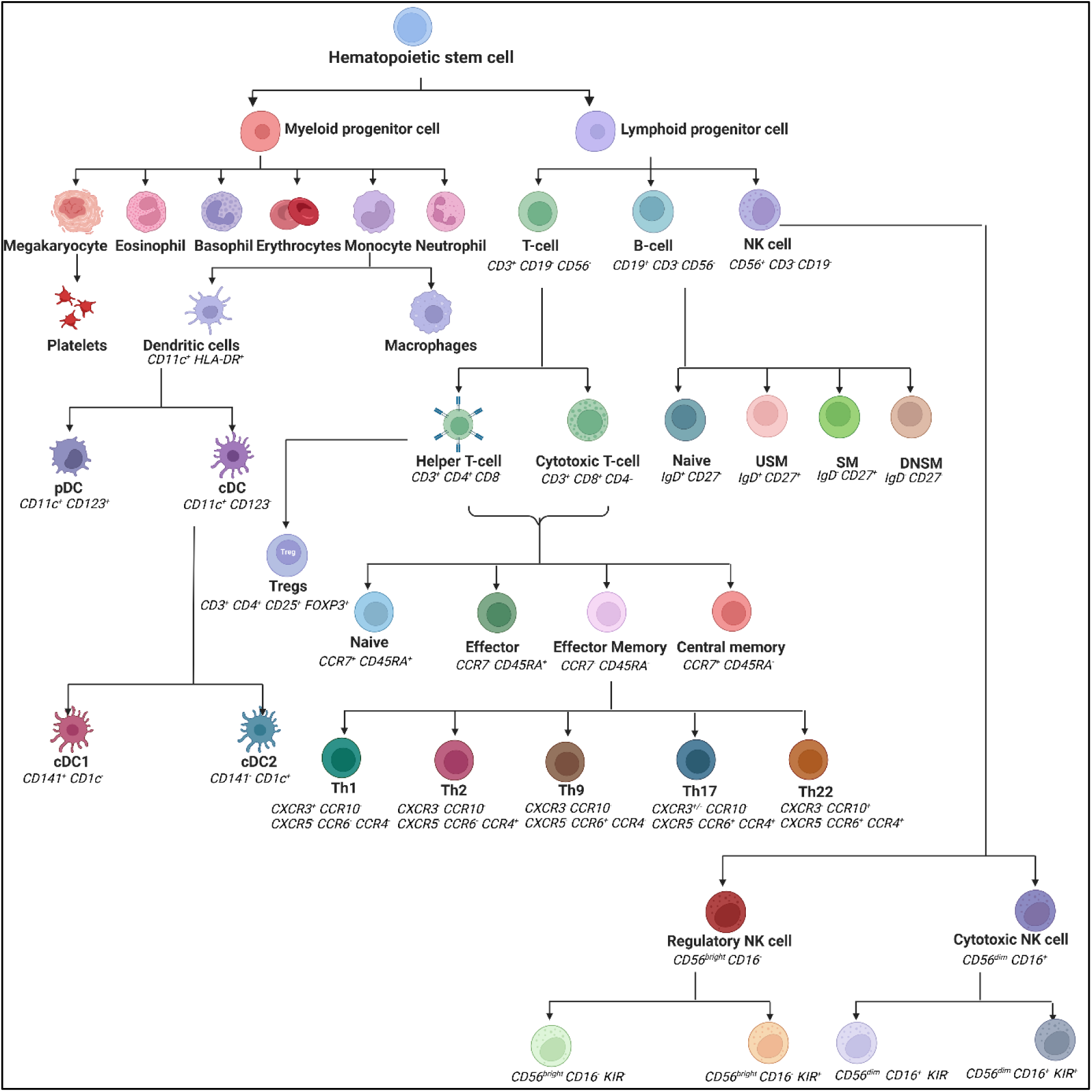
Schematic representation of immune cells and their subtypes, with profiling markers (mentioned in italics) used in the thesis for immunophenotyping. Abbreviations: NK: Natural Killer, pDC: Plasmacytoid Dendritic Cell; cDC: Conventional Dendritic Cell; Th: Helper T-cell; Tregs: Regulatory helper T-cell; USM: Unswitched Memory; SM: Switched Memory; DNSM: Double Negative Switched Memory

**Figure S2:**
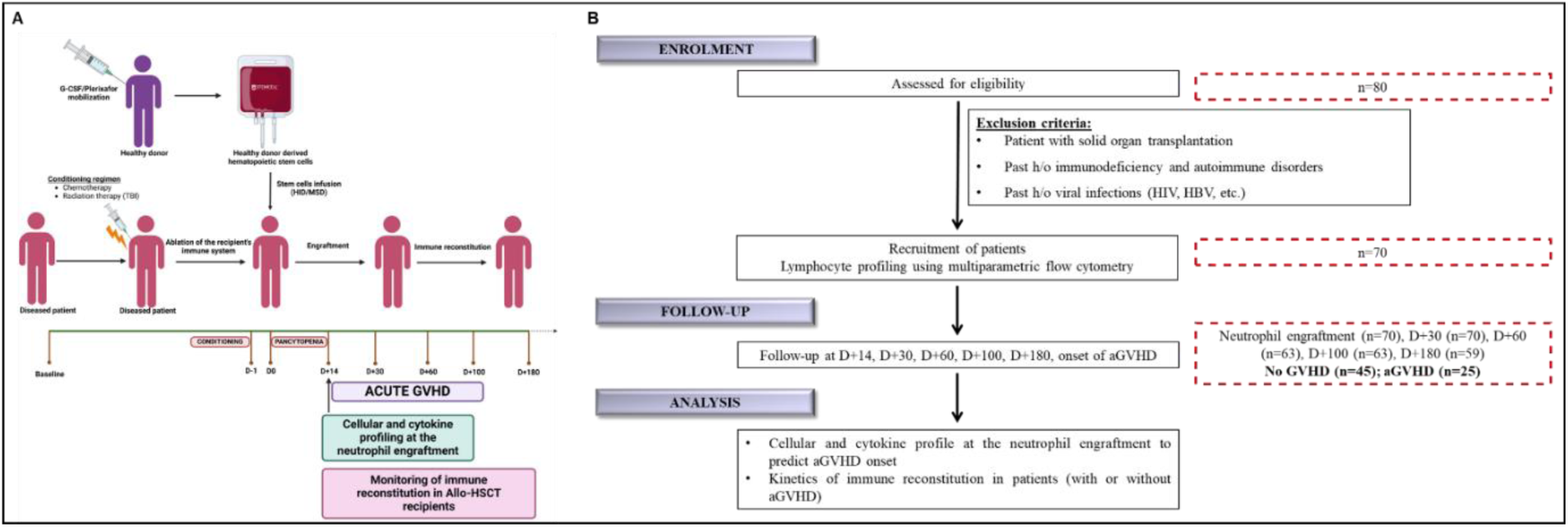
Schematic representation of (A) Study design. (B) Patient recruitment diagram of the study.

**Table S1:** Patient characteristics.

|  |  |
| --- | --- |
| **Age in years, median** |  |
| Patient | 21 |
| **Sex** |  |
| Male | 48 |
| Female | 22 |
| **Diagnosis** |  |
| AML | 25 |
| ALL | 8 |
| CLL | 1 |
| CML-blast crisis | 2 |
| Thalassemia | 9 |
| Aplastic Anemia | 16 |
| MDS | 3 |
| Others | 6 |
| **Donor** |  |
| Haploidentical | 21 |
| Matched sibling donor | 49 |
| **Conditioning regimen** |  |
| ATG only | 33 |
| ATG+PTCy | 1 |
| PTCy only | 3 |
| Others | 33 |
| **Total body irradiation** |  |
| Yes | 1 |
| No | 69 |
| **GVHD (grade II-IV)** | 25 |
| **GVHD prophylaxis** |  |
| CNI+MTX+/-Steroids | 47 |
| CNI+MMF+/-Steroids | 23 |

